# An automated workflow for multi-omics screening of microbial model organisms

**DOI:** 10.1101/2022.07.18.500181

**Authors:** Stefano Donati, Matthias Mattanovich, Pernille Hjort, Simo Abdessamad Baallal Jacobsen, Sarah Dina Blomquist, Drude Mangaard, Nicolas Gurdo, Felix Pacheco Pastor, Jérôme Maury, Rene Hanke, Markus J. Herrgård, Tune Wulff, Tadas Jakočiūnas, Lars Keld Nielsen, Douglas McCloskey

## Abstract

Multi-omics datasets are becoming of key importance to drive discovery in fundamental research as much as generating knowledge for applied biotechnology. However, the construction of such large datasets is usually time-consuming and expensive. Automation is needed to overcome these issues by streamlining workflows from sample generation to data analysis. Here, we describe the construction of a complex workflow for the generation of high-throughput microbial multi-omics datasets. The workflow comprises a custom-built platform for automated cultivation and sampling of microbes, sample preparation protocols, analytical methods for sample analysis and automated scripts for raw data processing. We demonstrate possibilities and limitations of such workflow in generating data for three biotechnologically relevant model organisms, namely *Escherichia coli*, *Saccharomyces cerevisiae*, and *Pseudomonas putida*.

## 1.1 Introduction

Microorganisms have evolved to live and thrive in a diverse set of environments ^123^. One of the factors that has contributed to this adaptability is the evolution and plasticity of their metabolism. Nowadays, manipulation of metabolism in microorganisms can be exploited for industrial biotechnology ^4^ as well as for biomedical purposes ^5^. However, current efforts to harness and engineer microorganisms for biotechnology and human health applications are limited by our understanding of biology. For example, in *Escherichia coli*, widely regarded as the best-studied microbial model organism, functional annotation of 34.6% genes is missing ^6^. Metabolite-protein interactions are also widely unknown ^7^ and only recently starting to be systematically mapped ^8–10^. In order to bridge these gaps in our understanding of biology at the systems level, a new wave of technological advances in laboratory automation, analytical chemistry, and data science is needed. In particular, automated cultivation platforms that can reproducibly grow a diverse range of microorganisms in different environmental conditions (e.g., degrees of aerobicity and ranges of temperatures) that support multi-omics (e.g., genomic, transcriptomics, proteomic, metabolomics, among others) data generation and analysis workflows are required ^11^.

Recent efforts have been made towards developing automated cultivation platforms that support a diverse range of cultivation conditions and volumes (See Ladner et al.^12^ for a review). Most analytical workflows for obtaining metabolomics, lipidomics, proteomics, and fluxomics data require several millions of cells per sample. Commercial automated cultivation platforms that fall into this range include 2Mag ^13–15^ and BioLector ^16–18^. The 2Mag reaction block supports 48 8-12 mL continuously stirred cultures grown aerobically or anaerobically with online pH and optical density (OD) sensors. The BioLector was designed to operate 48 well microtiter plates (MTPs) that can range from 800 to 2400 uL with online pH and dissolved oxygen (dO2) sensors and online OD measurements. Both the 2Mag reaction block and BioLector have been integrated with liquid handling robots to enable online pH control, substrate feeding, and sampling for other online or offline measurements ^13–18^. While both commercial 2Mag and BioLector platforms provide tight control of physiological conditions such as temperature, aeration, etc., several major limitations exist that limit their use for high throughput omics experiments. Limitations include the need for expensive disposable reaction chambers, limited throughput (i.e., only up to 48 cultivations at a time) and lack of support for fast sampling techniques for metabolomics, where speed of sampling is critical for accurate data acquisition ^19^. Customized automated cultivation platforms can also support a diverse range of growth conditions and experiment types. For example, a Tecan liquid handling robot was customized to support unicellular phototrophic growth in MTPs with control of CO_2_ and light ^20^. In another case, a Tecan robot was integrated with a customized 2Mag block with re-usable cultivation tubes and an OD reader for high throughput adaptive laboratory evolution experiments (ALE)^21^. In another example, a Hamilton robot was integrated with an incubator for growth in standard 96 well MTPs, an OD reader, and online analytical assays for pH, acetate, and glucose concentrations ^22^. A more recent approach involved a do-it-yourself (DIY) robotics system based on open-source components including Raspberry Pi, Arduino, and python that was shown to scale to several dozens of 40 mL glass cultivation chambers ^23^. Importantly many of these automation platforms are built for specific experimental set-ups (e.g., ALE), and none of these platforms support both aerobic and anaerobic growth, greater than 96 simultaneous cultivations, and fast sampling and quenching for omics sampling on the same robot out of the box.

In this work, we describe the establishment of a complex workflow for high-throughput multi-omics screening of microbial organisms (Figure 1A). The workflow is composed of an automated cultivation and sampling Tecan platform, Agilent Bravo sample handling robots for sample preparation, the relative analytical methods and data analysis pipelines. The Tecan cultivation platform (TCP) robot, thanks to a custom 3D printed lid for 96well plates allows to i) track growth of aerobic and anaerobic cultivations ii) obtain samples for omics analysis such as endo/exo metabolomics, proteomics and proteinogenic amino acid analysis. We demonstrate that the platform is capable of generating omics data for the biotechnologically relevant microbial model organisms *Escherichia coli, Saccharomyces cerevisiae*, and *Pseudomonas putida*.

**Figure 1:**
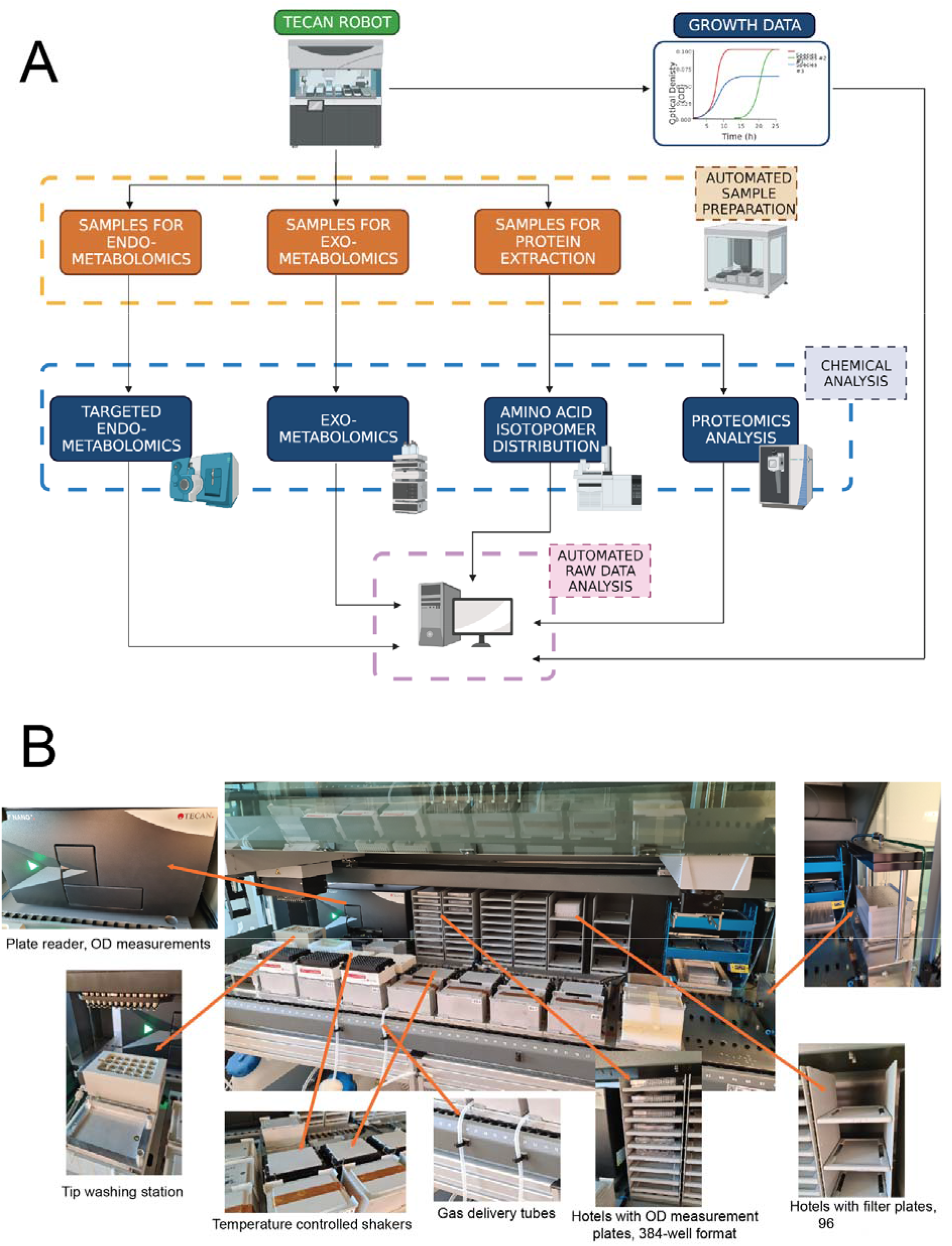
A) diagram representing the complete workflow. Cells were grown on the TCP and growth data was acquired using a plate reader. Upon triggering a user-defined OD, the cultivation platform could simultaneously sample for endo/exo-metabolomics or sample for protein extraction. Samples were then prepared and extracted for analysis using different workflows on Agilent Bravo robots. Processed samples were analyzed with the appropriate analytical instrument. All instruments were equipped with a 96-well format autosampler. Raw data was then processed through automated workflows. B) overview of the custom built TCP platform and its key components.

## 2. Material and methods

### 2.1. Design of a Tecan platform for automated and high throughput growth screening and sampling

A Tecan Freedom Evo liquid handler formed the base of the TCP (Figure 1B). The platform was integrated with the following main components: a Tecan Nano Plus spectrophotometer with adapters for online measurements at OD600 (culture biomass) and OD450 (OrangeG for volume loss control); a water reservoir for diluting samples for OD measurements; an online positive pressure station (Tecan Resolvex M10) for biomass separation from culture prior to quenching; plate hotels for 384-well plates (for OD measurement) and filter/collection plates (for biomass/culture separation); shakers (Bioshake 3000-T elm shaker, Qinstruments) for shaking and temperature control of the cultivation plates. Furthermore, tip boxes were dedicated for culture sampling (1-box of tips per cultivation plate) and diluting samples for OD measurement (1-box of tips). Several iterations of the deck layout were explored before settling on the current layout which was found to maximize cultivation capacity along with on deck tips and plate storage. With the chosen deck setup, the automated platform could perform OD measurements of a whole 96-well plate approximately every 8 minutes. The simultaneous operation of six 96well-plate cultivations with online OD sampling and OD triggered omics sampling was orchestrated by a script written using the Tecan EvoWare software with integrated Visual Basic modules for more complicated calculations. The main operations of the platform can be divided into two parts: 1) growth phase and 2) sampling phase (Supplemental Video 1). All scripts and user input files for running the TCP can be obtained by request.

During the fully automated growth phase, 5uL of culture was sampled from each cultivation plate and every well at predefined time intervals to measure culture density. Those 5 uL of culture were then diluted 5 fold (final volume 25uL) with filtered water in one of the quadrants of a 384-well plate (OD measurement plate) before OD measurement by the on-deck spectrophotometer. OD was measured at both 450 and 600 nm. Sampling tips were then washed using ethanol and water in the washing station and returned to their dedicated positions on the deck. After exhausting all four quadrants of the 384-well OD measurement plate, the spent plate was moved to trash and replaced by the next available 384-plate stored in the hotels.

The TCP allows for whole-plate semi-automated sampling for omics data generation. Upon reaching a particular cell density in pre-defined wells, cultures were processed in different ways depending on the desired type of sample and relative omics analysis to be performed. If the culture was used for endo/exo- metabolomics sample preparation, the culture was pipetted from the cultivation plate and moved to a filter/collection plate, which was subsequently filtered by positive pressure filtration. During positive pressure filtration the biomass was separated and maintained on the filter plate (for endo-metabolomics) and the supernatant deposited in the collection plate (for exo-metabolomics). After filtration, the filter plate was immediately moved (manually) and quenched by flash freezing the biomass in liquid nitrogen, and then both collection and filter plates stored in −80°C until further sample extraction. If the cultures were used for proteomics or fluxomics sample preparation, then the cultivation plates were moved off the deck of the TCP and centrifuged to pellet the biomass for storage in −80°C until further sample extraction.

### 2.2. Chemicals and reagents

Uniformly labeled ^13^C and 1-^13^C glucose was purchased from Cambridge Isotope Laboratories, Inc. 1,2-^13^C and 1,6-^13^C glucose along with unlabeled media components were purchased from Sigma-Aldrich-Merck. LC-MS reagents were purchased from Fisher Scientific and Sigma-Aldrich-Merck. Cultivation and sample handling materials were purchased from Waters and Sigma-Aldrich-Merck.

### 2.3. Organisms and cultivation conditions

*Escherichia coli* K-12 MG1655 was grown in glucose M9 minimal media. Glucose M9 minimal media consisted of 4 g/L glucose, 0.1 mM CaCl_2_, 2.0 mM MgSO_4_, trace element solution, and M9 salts. A 4,000× trace element solution contains 27 g/L FeCl_3_·6H_2_O, 2 g/L ZnCl_2_·4H_2_O, 2 g/L CoCl_2_·6H_2_O, 2 g/L NaMoO_4_·2H_2_O, 1 g/L CaCl_2_·H_2_O, 1.3 g/L CuCl_2_·6H_2_O, 0.5 g/L H_3_BO_3_, and concentrated HCl dissolved in double-distilled H_2_O (ddH_2_O) and sterile filtered. A 10× M9 salts solution contains 68 g/L Na_2_HPO4 anhydrous, 30 g/L KH_2_PO4, 5 g/L NaCl, and 10 g/L NH_4_Cl dissolved in ddH_2_O and autoclaved. *Saccharomyces cerevisiae* CEN.PK113-7D was grown in Delft media supplemented with 20g/L glucose^24^. *Pseudomonas putida* KT2440 was grown in de Bont medium^25^ containing 4 g/L of glucose as sole carbon source, 1.55 g/L K_2_HPO_4_, 0.85/L g NaH_2_PO_4_, 2.0 g/L (NH_4_)_2_SO_4_, 0.1 g/L MgCl_2_, 10 mg/L EDTA, 2 mg/L ZnSO_4_, 1 mg/L CaCl_2_, 5 mg/L FeSO_4_, 0.2 mg/L Na_2_MoO_4_, 0.2 mg/L CuSO_4_, 0.4 mg/L CoCl_2_, and 1 mg/L MnCl_2_. Obligate anaerobic bacteria were cultivated on modified GAM (mGAM) media.

### 2.4. Custom 3D printed lid

The lid was designed using SolidWorks CAD software and the model files are provided in the Supplemental Material. The three parts of the lid were 3D-printed using Stratasys Fortus 380mc printer from biocompatible Acrylonitrile butadiene styrene (ABS). The parts were assembled to a functional lid by mounting the parts together and sealing them with acetone to avoid any gas leakage between the parts.

### 2.5 Automated omics sample extraction and preparation

Automated cultivation and omics sampling was supplemented with automated omics sample extraction and analytical sample preparation implemented on Agilent Bravo liquid handling robots. Sample preparation methods included polar metabolite extraction for acquisition by liquid chromatography tandem mass spectrometry (LC-MS/MS), proteinogenic amino acid extraction and derivatization for acquisition by gas chromatography mass spectrometry (GC-MS) and proteomics sample preparation and acquisition by LC-MS. Other automated analytical workflows included the following procedures: dilution series of standards for calibration curves, plate replication, sample pooling and specific dilution of individual samples from a microtiter plate.

#### 2.5.1 Endo-metabolomics extraction and preparation

Endo-metabolomics samples from *E. coli* and *P. putida* were prepared using several automated steps: 1) the biomass was collected by positive pressure filtration (Tecan Resolvex M10) using Sirocco protein precipitation filter plates (Waters) on the deck of the TCP; 2) the filter plate was immediately removed from the deck of TCP and immersed in liquid nitrogen to quench cellular metabolism, and subsequently stored in −80°C for further extraction. All further liquid handling steps except filtration were done by the Agilent Bravo: 1) metabolites from the filter plate were extracted by adding 150μL of cold (−20 °C) extraction solvent (ES: 40% (v/v) acetonitrile (ACN), 40% (v/v) methanol and 0.1 M formic acid) and internal standard (IS, 13C labeled). Internal standards for metabolomics were produced by growing *E. coli* on M9 minimal media containing labeled glucose and extracting the labeled metabolomes. After the addition of ES and IS, the filter plate was immediately incubated for 2 hours at −20 °C; 2) the filter plate was filtered by positive pressure and washed twice with 150μL of cold (−20 °C) ES by filtration after each wash; 3) collected filtrate was evaporated dry over-night in a concentrator (Eppendorf^®^ concentrator plus) under vacuum conditions; 4) dried samples were reconstituted in 100μL LC-MS grade water (Fisher Chemicals) and transferred to a filter plate (AcroPre filter plate 0.2 μm (8019) - Pall) for further clean-up; 5) The filter plate was filtered by positive pressure and the collected flowthrough was ready to be injected into mass spectrometer. For *S. cerevisiae* samples an additional extraction step with boiling ethanol was performed ^26^. After the filter with biomass was stored at −80°C: 1) metabolites from the filter plate were extracted by adding 150μL of boiling (75-80°C) extraction solvent (ES: 70% EtOH) and internal standard (IS, 13C labeled). After addition of ES and IS the filter plate was immediately stored for 5 minutes at 70°C; 2) the filter plate was filtered by positive pressure and washed twice with 150μL of boiling (75-80°C) ES by filtration after each wash. The rest of the steps for sample preparation were the same as described above for the cold extraction procedure.

#### 2.5.2 Exo-metabolomics extraction and preparation

Exo-metabolomics samples from all three model organism were collected using the same protocol: 1) the biomass from the culture was separated by positive pressure filtration (Tecan Resolvex M10) using Sirocco protein precipitation filter plates (Waters) on the deck of the TCP; 2) the filter plate was removed, while the filtered flowthrough was kept, and subsequently stored in −80°C for further analyses by high performance liquid chromatography (HPLC).

#### 2.5.3 Proteomics sample extraction and preparation

Samples for proteomics were obtained by stopping the TCP run, removing the cultivation plate off the deck and collecting the biomass by centrifugation at 4000rpm for 5 minutes (not filtration as for endo- and exo-metabolomics), discarding the supernatant. Samples were kept in plates at −80 °C until processing, which was carried out in 96 well format. After samples were thawed on ice, two 3-mm zirconium oxide beads (Glen Mills, NJ, USA) and 100 μl of 95°C GuanidiniumHCl (6 M Guanidinium hydrochloride (GuHCl), 5 mM tris(2-carboxyethyl)phosphine (TCEP), 10 mM chloroacetamide (CAA), 100 mM Tris–HCl pH 8.5) was added to all the samples. Cells were disrupted in a Mixer Mill (MM 400 Retsch, Haan, Germany) set at 25 Hz for 5 min at room temperature, followed by 10 min in a thermo mixer at 95°C at 600 rpm. Any remaining cell debris was removed by centrifugation at 5000g for 10 min, after which 50 μl of supernatant was collected and diluted with 50 μl of 50 mM ammonium bicarbonate. Based on protein concentration measurements (BSA), aliquots with 100 μg of protein extract were used for tryptic digestion. Tryptic digestion was carried out at constant shaking (400 rpm) for 8 h, after which 10 μl of 10% trifluoroacetic acid (TFA) were added. Finally, samples were de-salted using SOLAμ C18 plates (Thermo).

#### 2.5.4 Amino acid isotopomer analysis sample extraction and preparation

Samples for amino acid isotopomer analysis were obtained in the same way as for proteomics analysis. After centrifugation the supernatant was removed and cell pellets stored in −80°C. Samples were extracted and prepared using the following procedure: 1) hydrolysis was performed by resuspending the cell pellets in 200μL of 6M HCl and transferring to 300μL - size FluidX tubes (Brooks Life Sciences). The tubes were sealed with FluidX rubber sealing mats (Brooks Life Sciences) and incubated for 12-16 hours in a heating block at 110°C; 2) after hydrolysis the lysates were moved to a filter plate (AcroPre filter plate 0.2 μm (8019) - Pall), filtered by positive pressure and the flowthrough was collected in a glass coated microtiter plate (WebSeal Plate+ 96-Well Glass-Coated, 300 - Fisher Scientific); 3) flowthrough was evaporated in the fume hood by heating samples at 60 °C with a thermomixer; 4) dried samples were derivatized on the Agilent Bravo robots, by adding 35μL of MTBSTFA (with 1% TBDMSCl) (Sigma-Aldrich) and 70μL of Pyridine (Sigma-Aldrich) to each well of the microtiter plate; the plate was then sealed and incubated at 65 °C for 30 minutes; 5) the samples were then transferred to HPLC glass vials for analysis.

### 2.6. Analytical chemistry methods

#### 2.6.1 Intracellular targeted metabolomics analysis

Intracellular metabolites were acquired and quantified on an AB SCIEX Qtrap® 5500 mass spectrometer (AB SCIEX, Framingham, MA) as described previously ^27^. Internal standards were generated as described previously ^28^. All samples and calibrators were spiked with the same amount of internal standard taken from the same batch of internal standards. Calibration curves were run before all biological and analytical replicates, and the consistency of quantification was checked by running a Quality Control sample that was composed of all biological replicates periodically between samples. Solvent blanks were injected periodically between samples to check for carryover. System suitability tests were injected daily to check instrument performance.

#### 2.6.2 Extracellular metabolomics analysis

Samples to determine substrate uptake and secretion rates were measured using refractive index (RI) detection by HPLC (Thermo Ultimate 3000) with a Bio-Rad Aminex HPX87-H ion exclusion column (injection volume, 10 ul) and 5 mM H2SO4 as the mobile phase (0.6 ml/min, 45°C).

#### 2.6.3 Proteomics measurement

For proteomics analysis a CapLC system (Thermo scientific) coupled to a Orbitrap Exploris 480 mass spectrometer (Thermo Scientific) was used. First samples were captured at a flow of 10 ul/min on a precolumn (μ-precolumn C18 PepMap 100, 5μm, 100Å) and then at a flow of 1.2 μl/min the peptides were separated on a 15 cm C18 easy spray column (PepMap RSLC C18 2μm, 100Å, 150 μmx15cm). The applied gradient went from 4% acetonitrile in water to 76% over a total of 60 minutes. While spraying the samples into the mass spectrometer the instrument operated in data-independent acquisition (DIA) mode using the following settings. The DIA method consisted of multiple MS1 scan with a DIA segment in between. The MS1-scan covered a range from 400-1250 m/z with the Orbitrap resolution set to 120 000; AGC Target 300%; maximum injection time Auto. The individual DIA segments consisted of 15 m/z windows at 30 000 resolution, normalized AGC target 1000% and injection time set to auto. The method in total lists like this: MS1 scan, DIA segment 400-670 m/z, MS1 scan DIA segment 400-670 m/z as previously described followed by a DIA segment 670-940 m/z, MS1 scan DIA segment 940-1210 m/z.

#### 2.6.4. Amino acid isotopomer analysis

Isotopomers of proteinogenic amino acids were acquired on an Agilent 5977 GC-MS system. Derivatized samples were run within 48 hours on the GC-MS, using an Agilent DB-5ms capillary column (30m, inner diameter of 0.25 mm, film thickness of 0.25 μm, cat no. 122-5532). Samples were measured in full-scan mode, using a 1:10 and 1:100 split ratio, with the following gradient: start at 160 °C, hold for 1 min, ramp to 310 °C at 20 °C/min, hold for 1 min. We considered for further analysis fragments listed in Table 3 of Long C.P. & Antoniewicz (2019) ^29^.

### 2.7. Data processing and analysis

#### 2.7.1. Growth data analysis

A custom Python-based module was developed to process the acquired OD measurements. This module combines all the individual 96-well plate OD spectrometer readings taken over the course of the experiment into a single spreadsheet, corrects volume loss due to dissipation and background noise, calculates the specific growth rates at each time point and best fit steady-state exponential growth rates for each well, and finally generates diagnostic figures. Volume loss over time is calculated by measuring the increase in intensity of OD450 measurements of a dye (OrangeG) in control wells in each cultivation plate. A linear model was fitted over the time series. Since volume loss causes the dye concentration to increase, the slope was expected to be positive. Correlation values were calculated for each timepoint by multiplying the slope with the timestamp and adding one to the intercept in order to leave the first time point unchanged. The experimental values were subsequently divided by the corresponding correlation value in order to decrease their adjusted concentration accordingly. The background correction was calculated by subtracting the mean of the OD600 measurements of the OrangeG control wells from the samples on the same cultivation plate for each time point. In addition to specific growth rate, the module used an in-house python package called croissance ^30^ to find the exponential growth phase and calculate the exponential growth rate. The quality control figures generated by the module included a visualization of the volume loss, the progression and distribution of the specific growth rates and distributions of initial values, the slopes and signal-to-noise ratios. The options for growth profiles included presenting the wells sorted by species and limited by a cultivation plate or vice versa, combined by species or cultivation plate or presenting the growth profiles of all the tested species combined in one plot.

#### 2.7.1. Metabolomics data analysis

Metabolomics raw data files were first converted to mzML format using ProteoWizard ^31^ and then processed using SmartPeak ^32^. Metabolite concentrations expressed in uM were normalized to cell biomass expressed in umol/gDCW at the time of sampling by interpolating the measured culture density during exponential growth to the expected culture density at the time of sampling and then compensating for the culture volume sampled and reconstitution volumes. The equations for biomass normalization were the following:

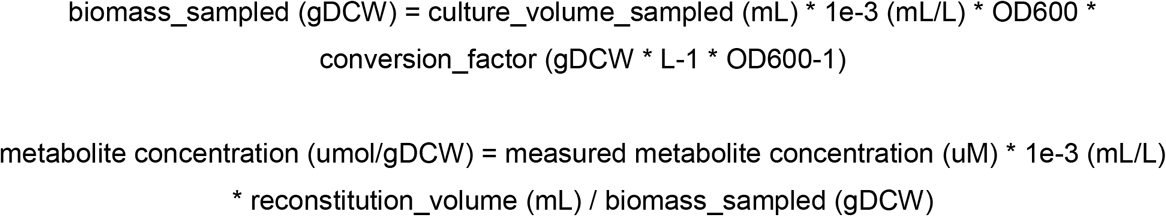

The energy charge ratio (EC) was calculated with the following equation:

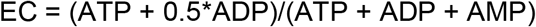

#### 2.7.2. Fluxomics data analysis

For analysis of amino acid isotopomer data, we considered fragments listed in Table 3 ^29^. Raw data from the GCMS was first converted to mzML format using ProteoWizard ^31^ and then processed using SmartPeak ^32^. Processed data was further corrected for the natural abundance of isotopes in the derivatization agents used for GCMS analysis ^33^ and analyzed with INCA ^34^.

#### 2.7.3 Phenomics data analysis

Extracellular metabolomics raw data in .txt format was processed using SmartPeak ^32^. Uptake and secretion rates were calculated from a minimum of three steady-state time-points taken from different cultivations at different dilutions. For calculations, we used the average maximum growth rate values reported in Table 1. We calculated the biomass concentration content at the time of sampling using OD/biomass conversion factors obtained using moisture analyzer (HC103 Moisture Analyzer - Mettler Toledo). The conversion factors used were the following: 2.06gDW/L/OD for *E. coli*, 3.62 gDW/L/OD for *P. putida* and 3.31 gDW/L/OD for *S. cerevisiae*. We considered only measurements for which the linear regression between the measurements at different dilutions had a squared correlation coefficient exceeding 0.8.

**Table 1:** maximum growth rates of microorganisms cultivated in the TCP compared to shake flasks. In the case of 96well growth rates, we calculated growth rates over time using a linear regression model with 3 measurements and reported the average maximum growth rate and its standard deviation across the whole plate (N=80). For flask cultivations, we report the average growth rate from 3 OD measurements and the relative standard deviation (N=3). Growth data is available in Supplementary Table 1.

| Organism | Growth rate measured plates | Growth rate measured flasks |
| --- | --- | --- |
| *E. coli* MG1655 (aerobic) | 0.63 +/- 0.06 | 0.61 +/- 0.05 |
| *E. coli* MG1655 (anaerobic) | 0.27 +/- 0.08 | 0.26 +/- 0.04 |
| *S. cerevisiae* CEN.PK122 | 0.44 +/- 0.08 | 0.33 +/- 0.07 |
| *P. putida* KT2440 | 1.09 +/- 0.06 | 0.7 +/- 0.09 |

2.7.4. Proteomics data analysis

Spectronaut 15.4 (Biognosys) was used to analyze the raw data files as direct DIA ^35^ with the following settings: Fixed modifications: Carbamidomethyl (C) and Varible modifications: oxidation of methionine residues; Trypsin as enzyme and allowing one missed cleavage; FDR set at 0.1%. Quantification was only based on MS1 intensities and normalization was set to global. For the searches, a protein database consisting of the reference proteomes UP000000625 (*E. coli*), UP000000556 (*P. putida*) and UP000002311 (*S. cerevisiae*).

## 3. Results

### 3.1. A custom made cultivation plate lid enables control of headspace gas for 96 well cultivations

Microorganisms have evolved to grow under various degrees of aerobicity. In order to cultivate such organisms, it is necessary to precisely control the headspace gas composition of their cultivation environment. To this end, we designed and fabricated a custom 96-well plate lid using 3D printing technology (Figure 2A). We used biocompatible Acrylonitrile butadiene styrene (ABS) which allows for chemical sealing between the layers of the lid to prevent air leakage and sterilization using ethanol and UV radiation. The lid was composed of three layers that are 3D-printed and chemically sealed using acetone. When assembled, the inner chamber of the lid formed a network of channels that aided in dispersing the air uniformly across all wells. The uniformity of airflow was assessed by submerging the lid under the water while pushing air through the lid to observe the density and distribution of bubbles that form. Automation friendly 96-well funneled sampling ports at the top of the lid allowed for robust and automated sampling during cultivation even when a portion of the sampling tips were offset. The airflow through the lid was such that air was pushed out of the sampling ports when not blocked by pipetting tips which prevented contamination and allowed for operation of the TCP without a HEPA filter and laminar flow.

**Figure 2:**
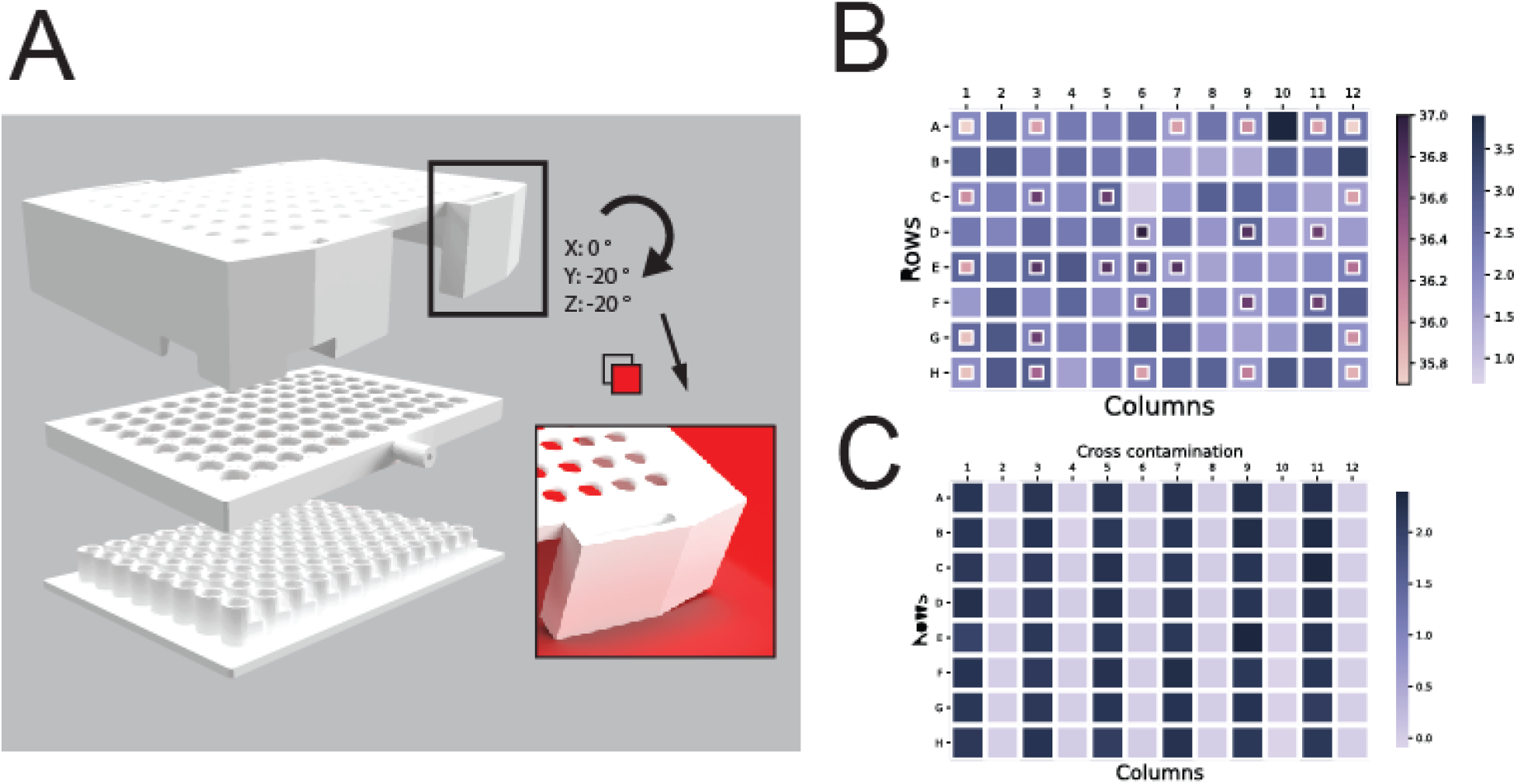
A) A custom lid was 3D printed to control the headspace gas of the cultivations. Consisting of 3 parts, the lid was assembled by stacking and chemical sealing. Once fixed on a cultivation plate, the created chamber allowed for controlling the headspace gas composition and evenly dispersed the air across the wells. The conical inlet holes for the sampling ports on the top of the lid are highlighted on the right. Background added for contrast. B) Temperature distribution and volume loss. The temperature of selected wells was recorded with a thermometer and is plotted in the small squares. The volume loss was estimated via the change of OD450 measurements of OrangeG and is plotted in the large squares. The temperature distribution indicates edge effects, the outside wells registering a lower temperature than the ones closer to the center of the plate. The volume loss does not appear to be correlated with the temperature distribution. C) All the wells of a 96-well microtiter plate were filled with media and every other column was inoculated with E. coli. OD measurements were recorded every 2 h. After 8 h, the growth rate for each well was calculated based on OD measurements. The inoculated wells showed growth rates in the expected range while no growth could be observed in the wells filled with only media.

96-well plate cultivations are notorious for edge effects, which can be broadly categorized as non-uniform heating and volume loss across all 96 wells. Edge effects contribute to high variance between replicates and non-reproducibility between experiments. In order to understand the extent to which edge effects were present when cultivating using custom 3D-printed lid, we quantified volume loss by measuring Orange G and temperature, using a thermometer (Figure 2B). Overall, the expected lower temperatures at the edges of the plate did not match the random distribution of evaporation rates, suggesting that the major factor in evaporation might be related to the air circulating in the lid.

Several measures were necessary to avoid cross contamination while conserving tips. First, 96-well plates were sealed with aluminum seal and pierced using a dedicated box of tips and the TCP MCA96 pipetting head prior to starting any cultivations. The aluminum seal was found to prevent cross talk between wells arising from condensation accumulation during cultivation. Second, a tip box was dedicated to each cultivation plate (i.e., 6 tip boxes for 6 cultivation plates). Third, tips were washed using ethanol and water in an on-deck washing station to remove culture components and disinfect tips following sampling for OD measurements. Cross contamination was evaluated by the following experiments: 1) *E. coli* MG1655 K-12 model organism was cultivated in plates where every other column was filled with only the media and assessed for growth after 8 hours (Figure 2C), and 2) a plate inoculated with a model organism and a plate filled only with media were sampled using the same tips for 24 hours and the plate filled with media was assessed for growth. In both experiments, no growth was observed in the wells filled only with medium, demonstrating that no cross-contamination was occurring either from improperly sterilized tips or from well-to-well in a cultivation plate.

### 3.2. Physiological characterization of model organisms in aerobic and anaerobic conditions

The ability of the TCP to ascertain accurate growth profiles was further tested using *E. coli, S. cerevisiae* and *P. putida* in aerobic conditions with minimal media (Figure 3A). The growth rates for *E. coli, S. cerevisiae* and *P. putida* were found to slightly differ from rates obtained in shake flask cultivations (Table 1), most likely due to differences in aeration conditions of the culture.

**Figure 3:**
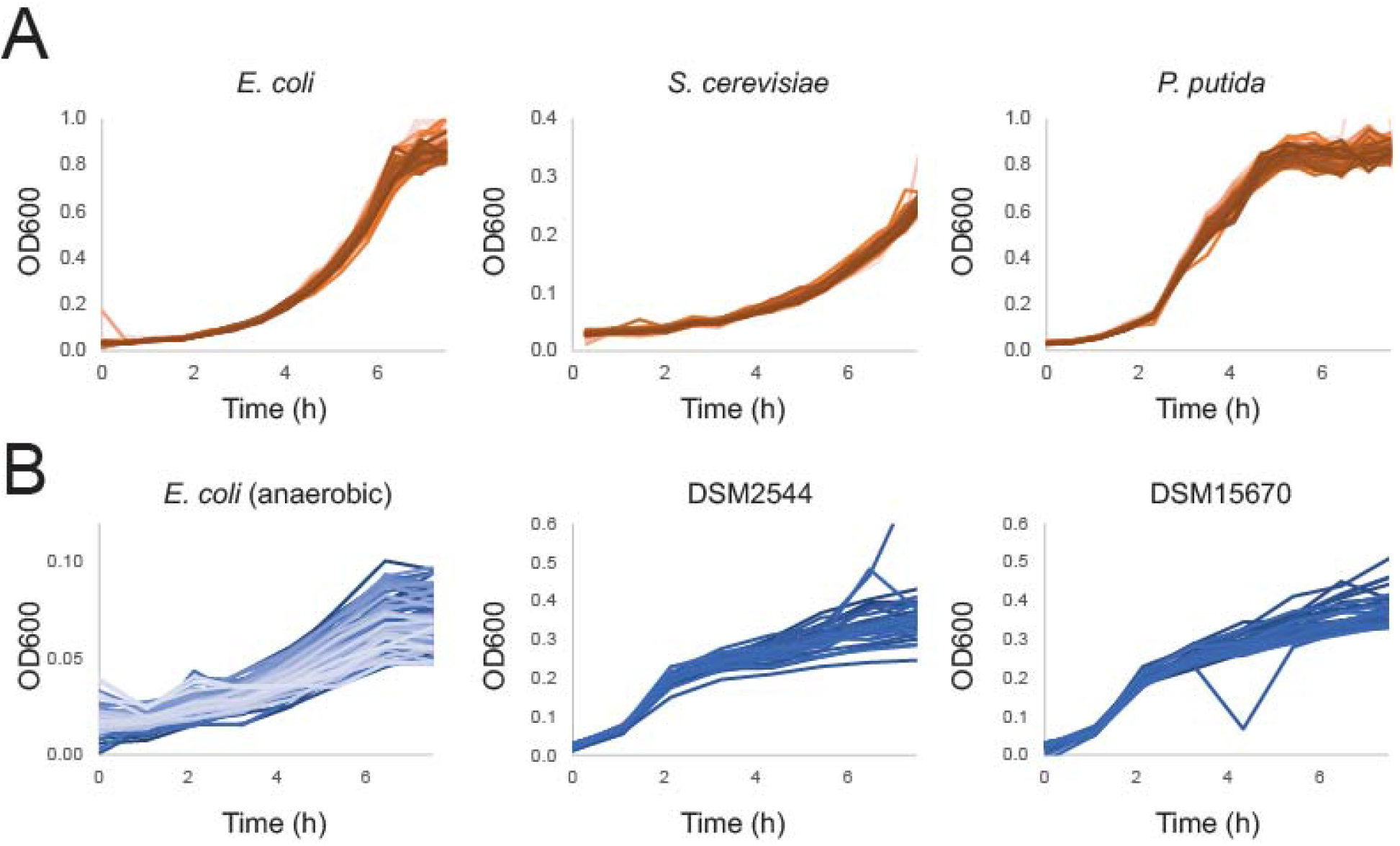
A) growth curves of aerobic cultivations on the TCP of 80 biological replicates of *E. coli, S. cerevisiae* and *P. putida*. B) growth curves of anaerobic cultivations of *E. coli* (N=40) and obligate anaerobs DSM2544 (*Lacrimispora saccharolytica*) and DSM15670 (*Enterocloster bolteae*).

Next, we tested whether the custom-made lid could be suitable for anaerobic cultivations. After preparing and sealing the cultivation plate in an anaerobic chamber, we covered the plate with the lid, flushing it with pure nitrogen instead of air. After the first OD measurement the aluminum seal was pierced, causing the liquid culture to be separated from the room atmosphere only by the nitrogen flushed through the lid. For *E. coli* in anaerobic conditions (Figure 3B, Table 1), we could observe lower growth rates than in aerobic conditions, as expected. We then tested the platform with two obligate anaerobic bacteria, *Lacrimispora saccharolytica* (DSM2544) and *Enterocloster bolteae* (DSM15670) (Figure 3B). In both cases we could observe growth, with circa four doublings over a few hours. Growth of strict anaerobic organisms confirms that the lid can assure anaerobic environment of the cultures.

### 3.3. A single time point strategy for capturing metabolic rates in plates

The ability to ascertain the uptake and secretion rates during steady-state growth enables several downstream analyses that are critical for quantitative modeling of organisms including constraint-based analysis (e.g., FBA, FVA, etc.) and absolute reaction rate determination using metabolic flux analysis (MFA). Traditionally, uptake and secretion rates are determined by measuring metabolites from multiple exo-metabolomic samples (Figure 4A) and growth over time during steady-state growth of the same culture. However, this approach is not amenable to high-throughput and low volume cultivation methods. Recently, it has been shown that samples taken from dilution series of microbial cultivations of *P. putida ^36^* could be used to quantify with confidence uptake and secretion rates (Figure 4B). We tested the same single-time-point strategy on *E. coli, P. putida* and *S. cerevisiae*, by collecting culture samples with the cultivation platform (Figure 4C). Samples were then measured with HPLC, to obtain concentrations of glucose and of organic acids produced by fermentation. We also took samples from batch shake-flask cultivations and compared the data.

**Figure 4:**
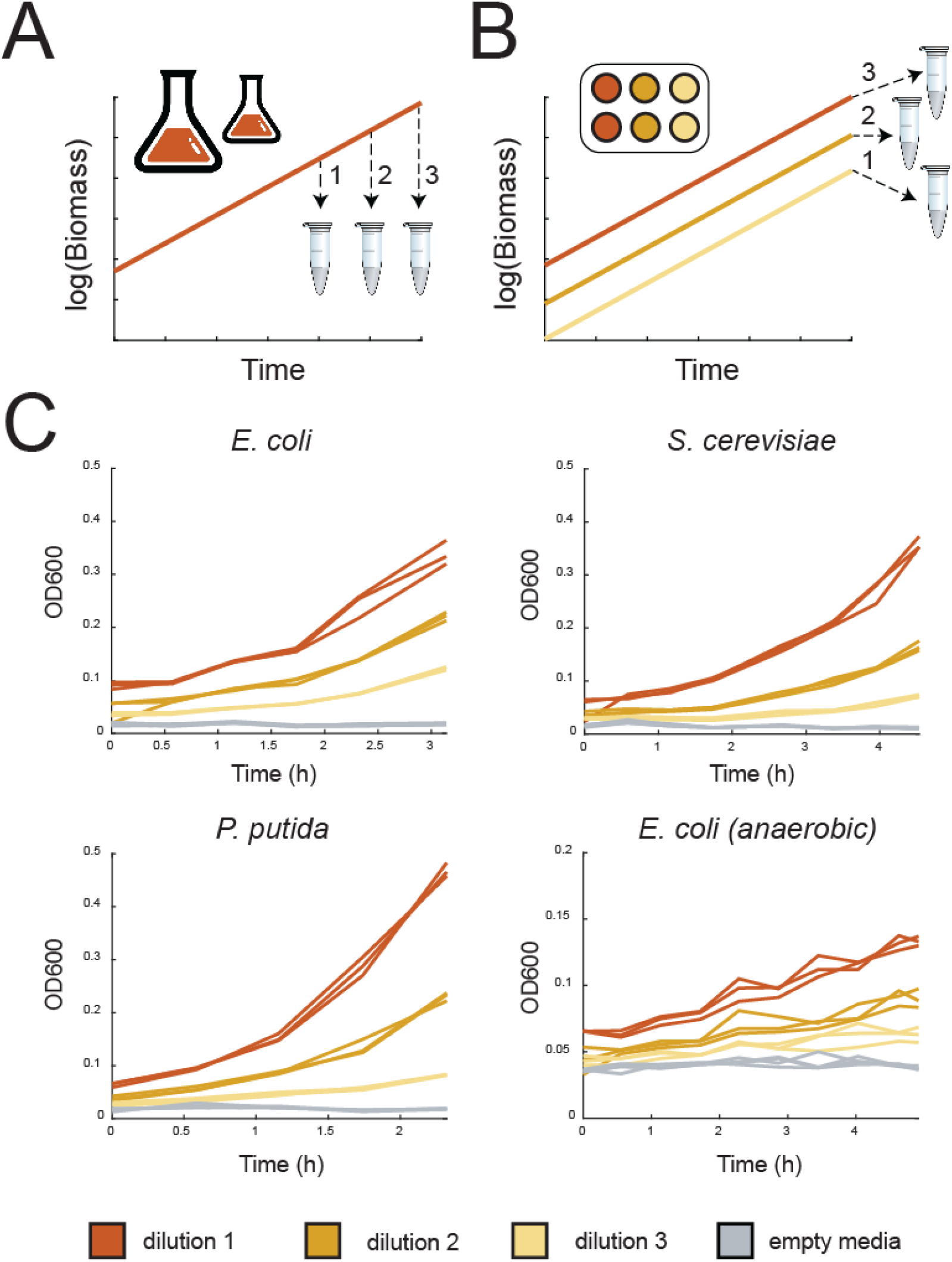
A) scheme of traditional sampling to calculate uptake/secretion rates B) scheme of single time point sampling strategy C) growth curves for all organisms of sampling experiments of the exo-metabolome. Samples collected at the end of the cultivation were used for HPLC analysis.

We observed that uptake and secretion rates obtained in the TCP could be quantified with a coefficient of variation lower than 30% among replicates. Aerobically, *E. coli* fermented mostly acetate, while anaerobically we observed mixed acid fermentation and strong accumulation of formate (Supplementary Table 2). However, formate formation in the aerobic cultivations of *E. coli* points to suboptimal aeration of the cultivations even at a measured growth rate greater than 0.6 1/h (Table 1). For *P. putida* we observed accumulation of gluconate and for *S. cerevisiae* strong production of ethanol. Overall, we show that the single time-point sampling strategy employed in our workflow produces meaningful data on the physiological state of the cell.

### 3.4. Integrated sampling and quenching for quantitative metabolomics captures a snapshot of cell physiology

The metabolome provides an instant readout of the cell state and requires rapid sampling and quenching protocols due to the fast turnover time of intracellular metabolites. Due to the diversity of metabolite chemical properties and the compositions of the cell wall and membrane, sampling, quenching, and extraction protocols are often compound-class and organism-specific ^37^. The metabolome is arguably the most sensitive cellular component, as it can dramatically change in a few seconds after perturbing cells ^38^. The required speed and need to accommodate heterogeneity make the automation of sampling and quenching for metabolomics is extremely challenging. An initial setup included automated quenching by fast addition of hot ethanol (70°C) on the filtered samples. However, LC-MS analysis of *E. coli* samples revealed a low energy charge ratio (EC) (Figure 5A). As samples were obtained from exponentially growing cultures, it was expected to observe an EC ratio close to 1 ^39^, and this discrepancy pointed to metabolite degradation during the quenching and extraction process. Hence, we set to test an acidic acetonitrile and methanol-based quenching solution ^40^ and compared it to the ethanol extraction method in various conditions, compatible with our automated 96-well format setup (Figure 5A, Supplementary Table 3). Eventually a semi-automatic procedure was chosen, which consisted in moving the filter plate with cells to a liquid nitrogen bath outside of the TCP, quickly incubating in −20°C and then subsequently adding cold ACN and incubating again at −20°C. The setup was chosen also as it allows to transfer the plate to a ventilated fume-hood ensuring safety measures when working with toxic volatile compounds such as acetonitrile. We then tested this protocol on a plate of *E. coli* cultivations. Samples were taken from exponentially growing cultures (Supplementary Figure 1) and measured with LC-MS. Overall, we could observe that the EC was evenly distributed throughout the plate (Figure 5B). While the EC was evenly distributed among the plate, absolute levels variated slightly (CV=<30%) if excluding 8 outlier samples out of 88 (Supplementary Figure 2). Similar results were obtained for whole plate cultivations of *P. putida* and *S. cerevisiae* (Figure 5C, Supplementary Figure 2). In summary, we could observe reproducible metabolome profiles among the whole plate and stable EC values, confirming previous findings that the custom manufactured cultivation lid is suitable to minimize edge effects and produce reproducible biological replicates. The sampling method proved to be applicable to *S. cerevisiae* and *P. putida* (Figure 5C). The few outliers present in the plates could be explained with clogging of a few wells on the filter plate during sampling and/or sample preparation.

**Figure 5:**
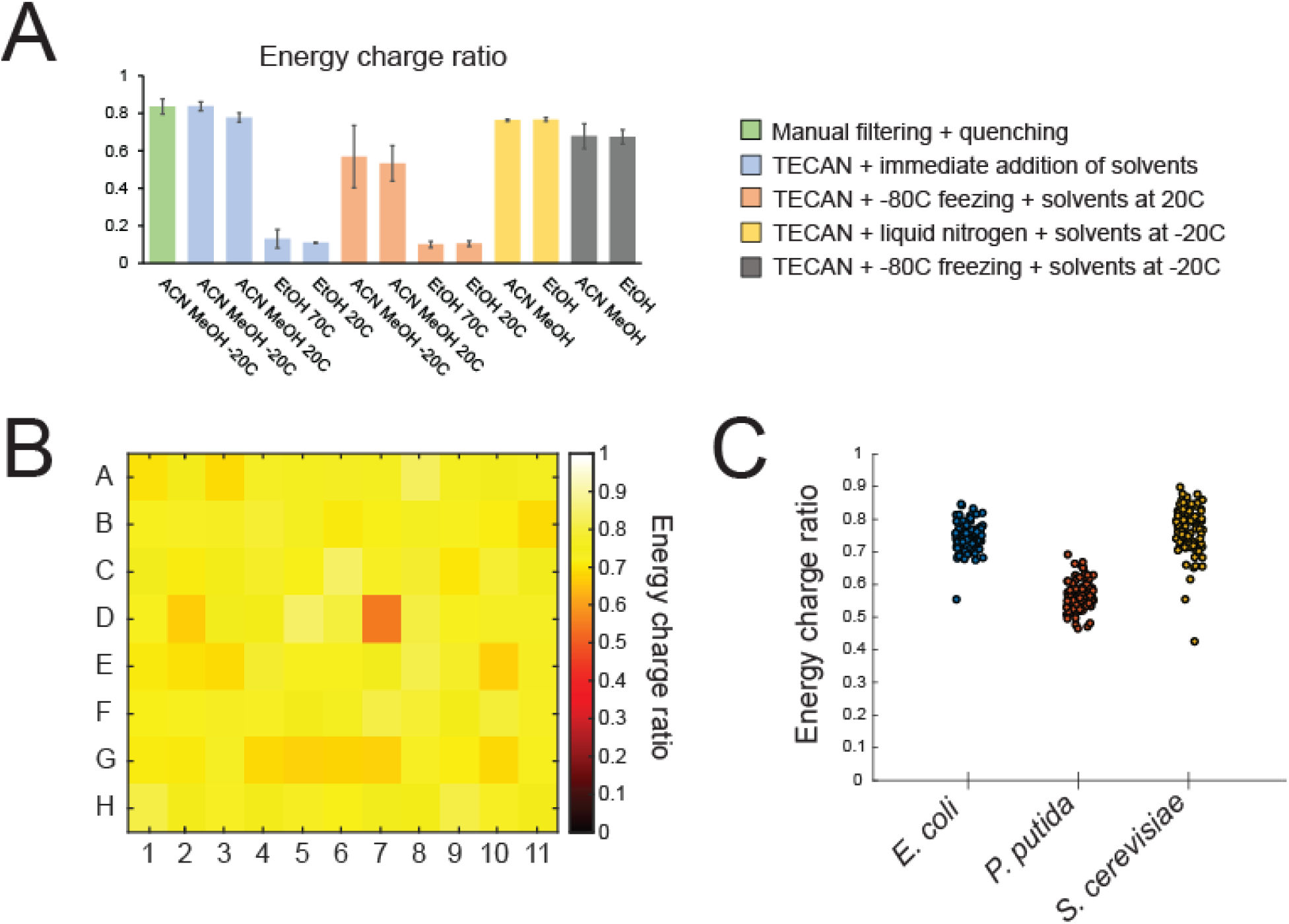
A) comparison of EC ratios among different quenching solvents and quenching conditions. Different colors represent the different conditions. Bars represent averages and error bars the standard deviations. B) heatmap displaying the distribution of the EC ratios among a plate of biological replicates of *E. coli*. C) distribution of values of EC ratios among 3 separate plates of cultivations of *E. coli, P. putida* or *S. cerevisiae* cultivations.

### 3.6. Proteome analysis of selected organisms

We verified edge effects of the plate-based cultivation and general quality of samples also at the proteome level. We cultivated separate plates with 80 replicates of *E. coli S. cerevisiae* and *P. putida*, observing again low variation in growth among replicates (Supplementary Figure 3) and obtaining biomass samples. After sampling at least 0.15 mg of biomass from each well. 21 randomly-selected samples, spread across the plate, were analyzed by DIA proteomics (Supplementary Table 4).

We observed a good reproducibility in coverage of the measured proteomes, with respectively 46%, 53% and 31% for *E. coli, P. putida and S. cerevisiae* of measured proteins out of protein coding genes. Moreover, most proteins displayed a low variability among replicates, with respectively 84% 81% and 85% of measured proteins retaining a CV lower than 30% among all replicates. Moreover, we did not observe strong outliers in the dataset when comparing random triplets of biological replicates (Supplementary Figure 4). This confirms that the proteome of biological replicates among the plates is reproducible and not affected by edge effects of the cultivation setup.

### 3.7. Parallel labeling experiments on the TCP

While the metabolome informs on concentrations of metabolic species, fluxomics methods allow to reconstruct the metabolic rates in the biochemical networks of organisms. An established method to reconstruct flux distributions is through 13C metabolic flux analysis (MFA) by measuring proteinogenic amino acids ^29,41^. This methodology involves sampling of cells grown on minimal media with labeled carbon compounds for isotopomer analysis of proteinogenic amino acids. This data in combination with metabolic uptake/secretion rates and growth rates, are then used to find the flux distribution that fits the data best in an assumed starting metabolic model.

We first measured mass distribution vectors (MDVs) of proteinogenic amino-acids of *E. coli* cultivated on glucose to assess the automated sample preparation pipeline and the accuracy of the GC-MS method. We compared our data to theoretical values calculated assuming natural isotope abundance of measured fragments (Supplementary Figure 5A). Overall, we observed a good precision of our measurements, with only a few fragments deviating by 1 mol% from the expected value. The ability of the TCP to obtain meaningful fluxomics data was then tested cultivating *E. coli* on different isotope tracers (1-2 13C glucose, 1-6 13C glucose) and collecting biomass samples in exponential phase. We then compared MDV data obtained from these experiments to similar experiments from literature, in which cells were cultivated in batch mode in small bioreactors ^42^. In this case, MIDs obtained from experiments on the TCP differed significantly, with transitions deviating up to almost 10 mol% from the reference dataset (Supplementary Figure 5B). Because of these differences, fitting the data on a reference metabolic model of *E. coli* did not result in an acceptable fit for MFA. Hence, it seems that in its current setup the TCP is not suitable for MFA experiments based on measurement of proteinogenic amino acids. This might be caused by difficulties in i) keeping cells in a controlled well-aerated environment ii) reaching isotopic steady-state.

## 4. Discussion

In recent years, multi-omics studies of metabolism have been broadening our understanding at the systems level, for example identifying novel metabolite-transcription factor interactions at a large scale ^8^, showcasing how cells can overcome mutations affecting metabolism ^43^ or perturbations of enzyme levels ^44^. The accumulation of a critical mass of high quality multi-omics datasets will eventually enable the application of machine learning methods to truly deconvolute biological complexity through the various biochemical layers that compose cells ^45^. In turn, this knowledge will allow the construction of more precise whole-cell models and as a consequence rational engineering of biological systems for biotechnological and medical applications.

We have described here the design and validation of a custom platform for automated and high throughput plate-based batch cultivation of aerobic and anaerobic microorganisms for omics analysis. We show that a custom 3D printed lid allows the control of headspace gas for plate-based cultivations, enabling for example anaerobic cultivations. Moreover, the integration of a positive pressure pump enables to effectively separate biomass from supernatant in an automated fashion. Effective sampling for metabolomics is a generally difficult task, due to instability of metabolites. We could show that combining sampling with filter plates and fast manual quenching in liquid nitrogen and cold organic solvents produces biologically relevant metabolomics samples, with only few outliers per plate. These outliers can potentially be caused by faulty filtration during sampling or sample preparation and/or uneven quenching of the plate in liquid nitrogen. Obtaining enough replicate samples (>3) from analyzed strains (which is trivial using the TCP described here) and discarding the faulty ones, based on analyzing growth profiles or filtration of the samples, would overcome this problem. Adapting the platform to operate safely with toxic organic solvents as acetonitrile, could also enable automated high-throughput quenching with possibly a lower number of outlier samples. Furthermore, we have shown that the platform is capable of taking highly reproducible samples for proteomics, extracellular metabolomics and amino-acid isotopomer analysis. Despite good data reproducibility, we observed some discrepancies from data generated in previous research studies. This discrepancy is most likely due to differences in aeration between 96 well plates against shake flasks and in the variability in reaching isotopic steady state in proteinogenic amino acids in short batch cultivations. Future work could explore performing MFA measuring metabolic intermediates ^46^ rather than proteinogenic amino acids.

The optimal plate layout for steady-state single time point omics sampling experiments consisted of 88 wells reserved for experimental conditions and 8 wells for controls such as growth and omics controls (e.g. *E. coli* MG1655 K12), contamination controls (empty media) and volume loss controls (OrangeG). We observed that the platform operates at best when cultivating 3 plates simultaneously, allowing growth measurements every 30 minutes. With such setup, the platform is capable of growing and sampling 264 batch cultivations per day. The capacity of this setup allows to generate in a few days hundreds of samples for multi-omics experiments, which would normally require many weeks of manual work by a researcher using traditional cultivation equipment (e.g., shake flasks). While online fluorescence measurements are already possible and routinely used, future work may look towards integrating other online measurements such as pH and dissolved oxygen, optimizing aeration conditions of the cultures, testing different headspace gasses and optimizing further quenching conditions for intracellular metabolomics.

## Supporting information

LID design

## 5. Acknowledgements

We would like to acknowledge Emre Özdemir for writing the parser translating the output of the Tecan OD reader to a summarized CSV file. Bo Hagelskjær Larsen from DTU Prototype Lab for 3D-printing the parts required to assemble the cultivation lid. Ricardo Hernández Medina for help in shooting the video of the TCP. This work was supported by The Novo Nordisk Foundation (NNF20CC0035580). Matthias Mattanovich was funded by the Novo Nordisk Foundation PhD fellowship (NNF19CC0035454).

## Author Contributions

S. D., P.H., S.A.B.J., N.G., J.M., R.H, T.J., D.Mc. participated in the development of the TCP. M.M., F.P.P. and D.Mc developed scripts for growth data analysis. S.D., S.D.B., R.H., J.M., and T.J developed sample preparation protocols. S.D., P.H., S.A.B.J., R.H., T.J. performed experiments. S.D., S.D.B, D.Ma., T.W. performed sample preparation, data acquisition and raw data processing. S.D. and M.M. analyzed the processed data. S.D. and D.Mc. wrote the manuscript. M.J.H. and D.Mc. designed the TCP L.N. and D.Mc. conceived of the study. M.J.H., L.N., and D.Mc. provided funding. S.D., M.J.H., T.J, L.N. and D.M. supervised the study.

