## Supplementary Figures for "An automated workflow for multi-omics screening of microbial model organisms"

Supplementary Materials

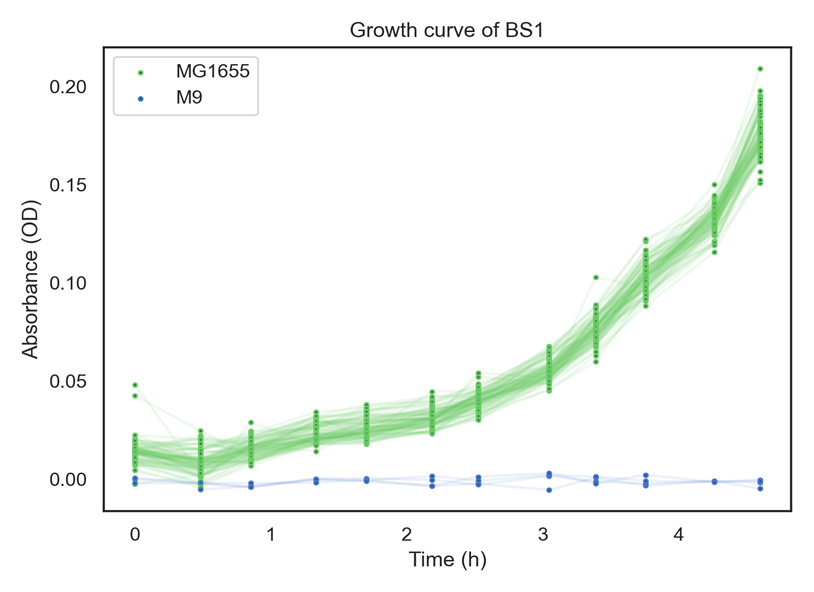

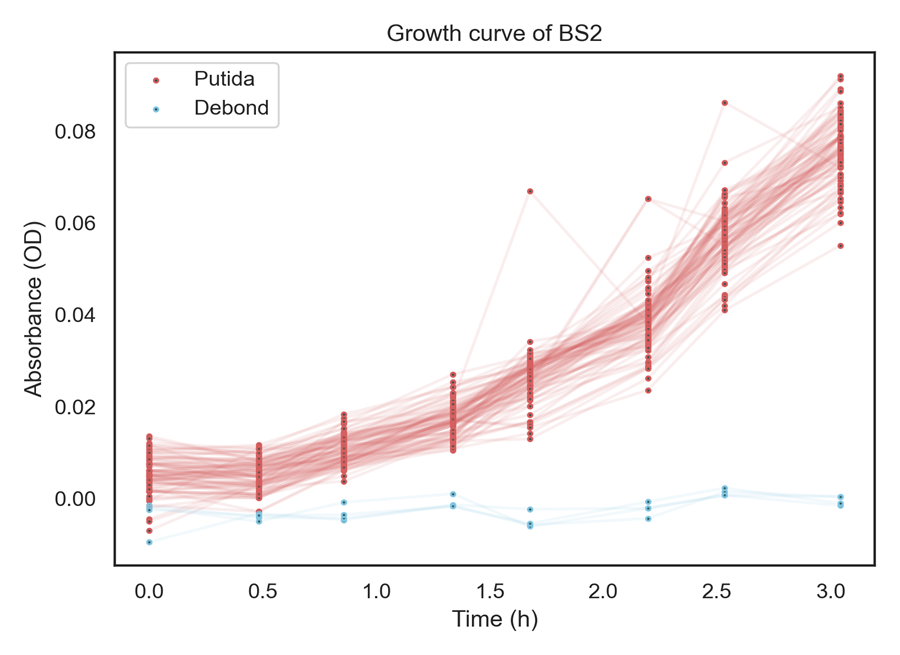

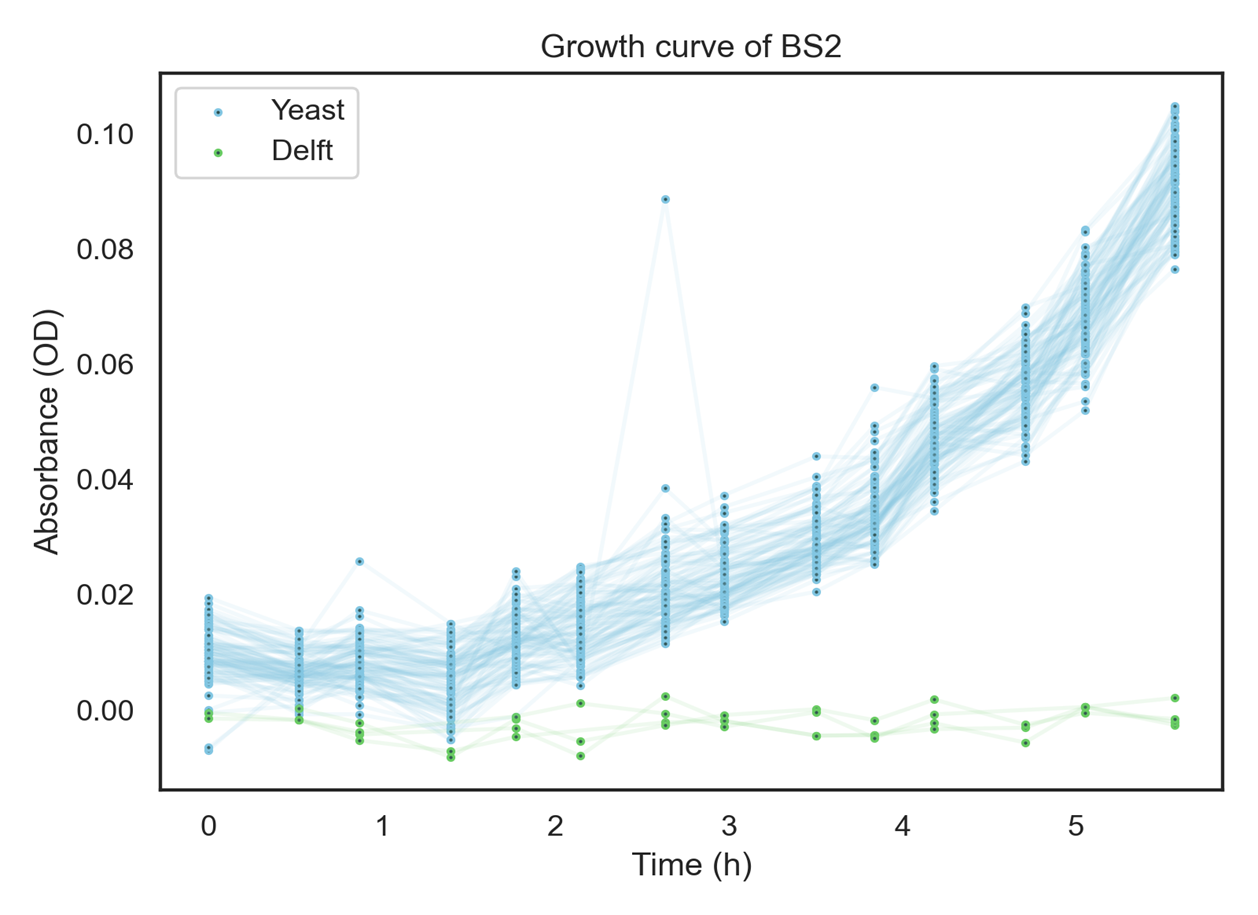

**Supplementary Figure 1:** growth curves of whole plate cultivations (n=80) before metabolomics sampling.

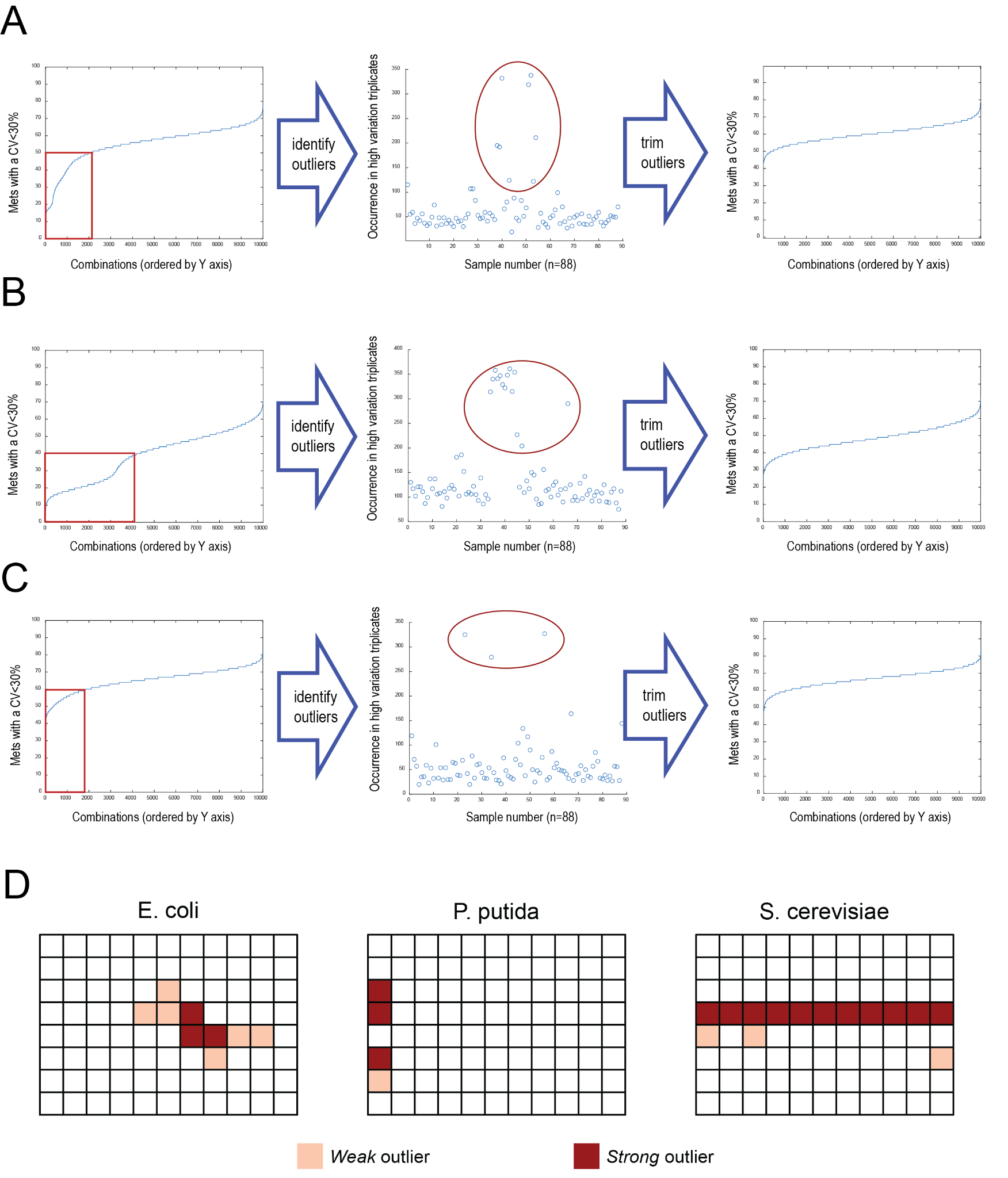

**Supplementary Figure 2:** variability of metabolomics samples and outliers A) number of measured metabolites in the *E. coli* metabolomics plate with a CV<30% among randomly generated triplets of sample data, ordered by the number of variability of metabolites. The occurrence of particular samples in the plate was identified, and upon removal of those outliers from the dataset the number of metabolites with a high CV decreased significantly. B) same as A, but for a plate of S. cerevisiae cultivations. C) same as A, but for a plate of P. putida cultivations D) distribution of the identified outliers over the plates. The row of outliers in the S. cerevisiae plate was caused due to a pipetting error when adding 13C internal standard to the samples.

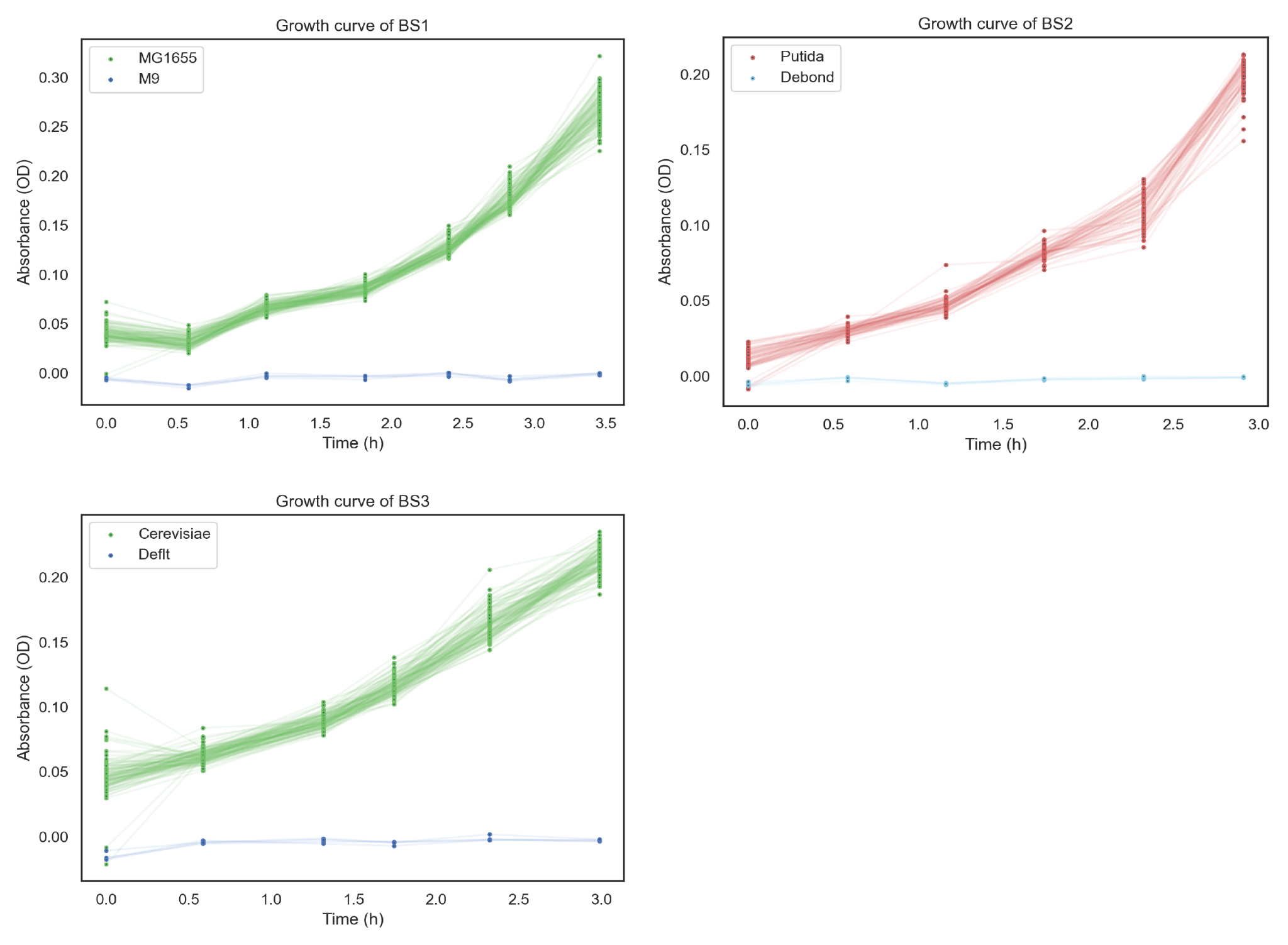

**Supplementary Figure 3:** growth curves of whole plate cultivations (n=80) before proteomics sampling.

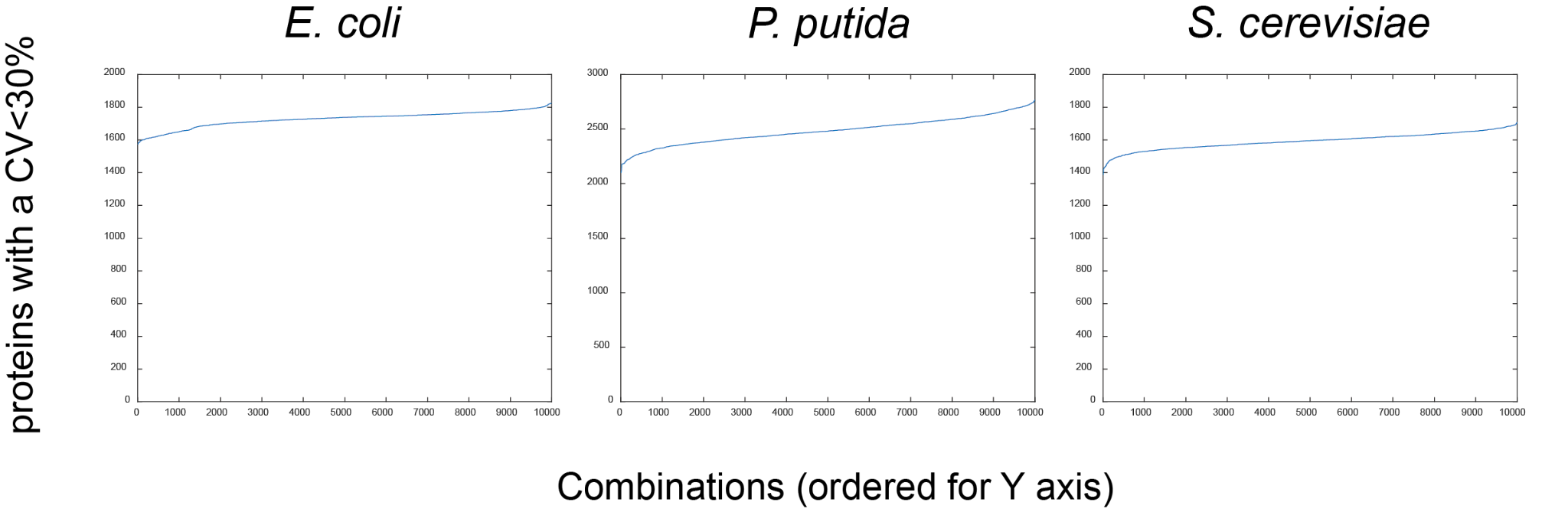

**Supplementary Figure 4:** number of measured proteins with a CV<30% among randomly generated triplets of sample data, ordered by the number of variability for the proteins, for the three analyzed organisms. The number of proteins with an acceptable CV is stable, regardless of the combination of sample data.

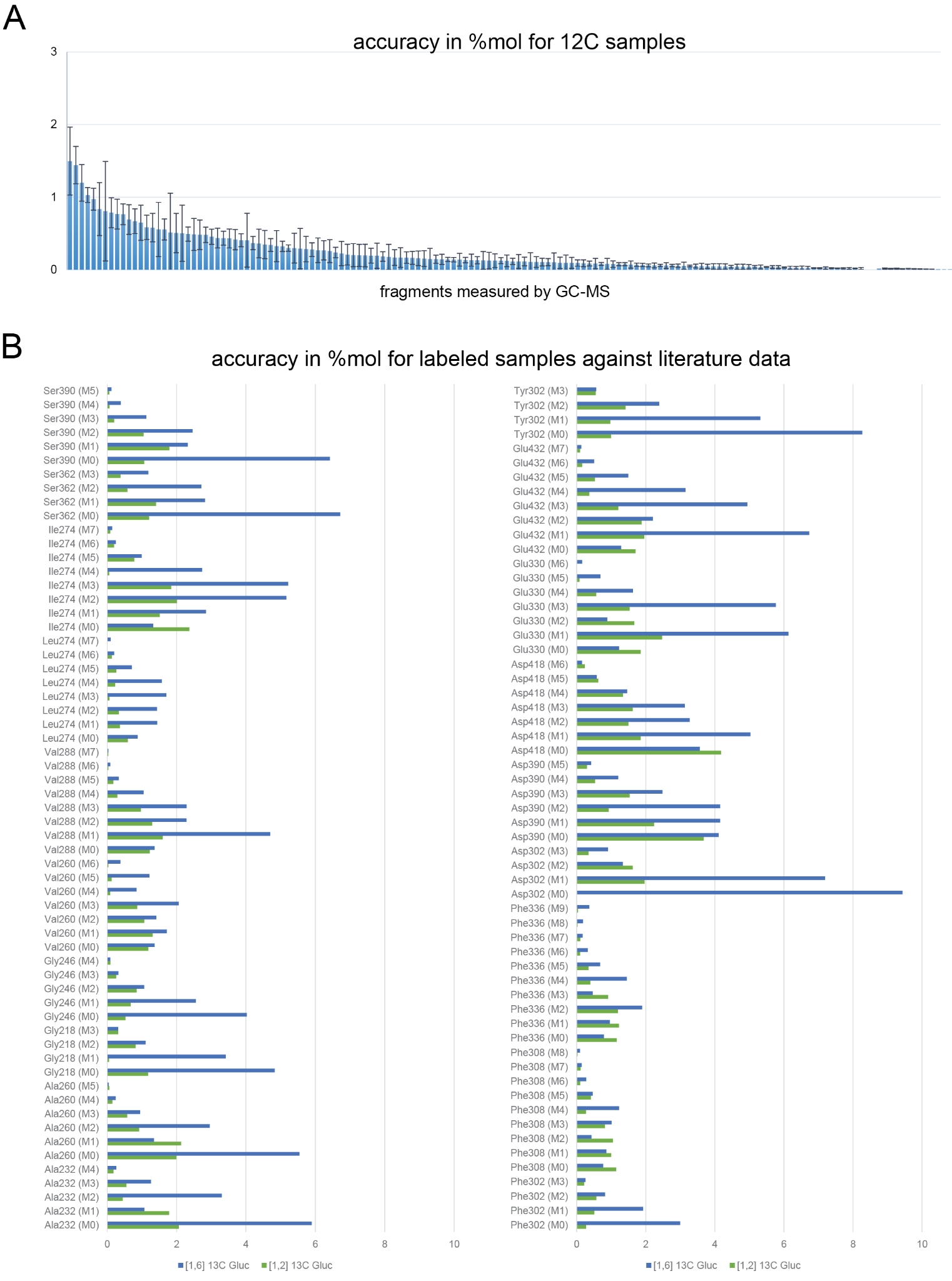

**Supplementary Figure 5:** accuracy of GC-MS measurements A) accuracy of the instrument for each measured fragment, quantified as the absolute value of the difference of measured MIDs of *E. coli* samples from cultivations on natural 12C glucose, against the theoretically calculated MIDs. Error bars indicate standard deviation of 8 replicates. B) accuracy of MIDs from experiments with labeled carbon sources (1,2 13C glucose and 1,6 13C glucose) on the TECAN in comparison with published data obtained in flask cultivations. Accuracy here is defined as the absolute value of the difference of measured MIDs from TECAN samples against the MIDs of samples from flask experiments.
